# Quantitative Models for Distinguishing Punctuated and Continuous-Time Models of Character Evolution and Their Implications for Macroevolutionary Theory

**DOI:** 10.1101/2024.04.09.588788

**Authors:** April M. Wright, Peter J. Wagner

## Abstract

The recent proliferation of quantitative models for assessing anatomical character evolution all assume that character change happens continuously through time. However, punctuated equilibrium model posits that character change should be coincide with cladogenetic events, and thus should be tied to origination rates. Rates of cladogenesis are important to quantitative phylogenetics, but typically only for establishing prior probabilities of phylogenetic topologies. Here, we modify existing character likelihood models to use the local cladogenesis rates from Bayesian analyses to generate the amounts of character change over time dependent on origination rates, as expected under the punctuated equilibrium model. In the case of strophomenoid brachiopods strop from the Ordovician, we find that Bayesian analyses strongly favor punctuated models over continuous-time models, with elevated rates of cladogenesis early in the clade’s history inducing frequencies of change despite constant rates of change per speciation event. This corroborates prior work proposing that the early burst in strophomenoid disparity reflects simply elevated speciation rates, *which in turn* has implications for seemingly unrelated macroevolutionary theory about whether early bursts reflect shifts in intrinsic constraints or empty ecospace. Future development of punctuated character evolution models should account for the full durations of species, which will provide a test of continuous change rates. Ultimately, continuous change vs. punctuated change should become part of phylogenetic paleobiology in the same way that other tests of character evolution currently are.

**Non-technical Summary:** Punctuated Equilibrium predicts a distribution of anatomical change that is fundamentally different from the models used in studies of relationships among species. We present a model to assess relationships that assumes punctuated change. We apply this model to a dataset of strophomenoid brachiopods to demonstrate that a model of punctuated change fits better than a model of continuous-time (“phyletic gradualism”) change in this group. Notably, because the punctuated model posits elevated speciation rates early in the strophomenoid history, the model also posits elevated rates of change among the early strophomenoids relative to later ones. This corroborates notions for what causes bursts of anatomical evolution rooted in ecological theory rather than evolutionary developmental theory. More basically, it emphasizes that paleontologists should consider both punctuated and continuous-time models when assessing relationships and other aspects of macroevolutionary theory.

## Introduction

Eldredge and Gould’s (1972) proposal that anatomical change commonly is “punctuated” rather than distributed continuously through time inspired a flurry of research. Although early research focused on particular case studies documenting examples of gradual (e.g., Gingerich 1979; Sheldon 1996) or punctuated (Cheetham 1986; Polly 1997) change, more recent studies have focused on the ubiquity of punctuation and stasis relative to continuous, anagenetic change (e.g., Hunt 2013; Hunt et al. 2015). Other work focused on the implications of punctuated change for the interface between macroevolution and processes such as selection (e.g., Lande 1976). However, the macroevolutionary implications of punctuated vs. continuous-time models go beyond how speciation itself works. This was first realized for trends over time (Eldredge and Gould 1972; Stanley 1975), as state-dependent origination and extinction rates (i.e., species selection *sensu lato*) can induce the same patterns as frequent shifts to particular conditions. The possibility of punctuated change has necessary implications for patterns of morphological evolution: punctuated change coupled with elevated cladogenesis rates should induce rapid increases in disparity and apparently high rates of anatomical change per million years even if the rates of change per speciation event remain constant (Foote 1996b; Congreve et al. 2021).

That macroevolutionary implications of punctuated change dovetail into methodological implications for how we conduct phylogenetic analyses with anatomical data. This goes beyond the co-occurrence of ancestor and descendant species that led to the initial recognition of punctuated equilibrium (Eldredge 1971) or the expected differences in tree-shapes that we expect from punctuated equilibrium and continuous-time models of character change (Wagner and Erwin 1995). Quantitative phylogenetic methods consider not only topologies, but also rates of change and divergence times (e.g., Lewis 2001). Rates of cladogenesis are important in Bayesian phylogenetic analyses: but to establish prior probabilities of divergence times and tree shapes (e.g., Heath et al. 2014), not for evaluating character evolution. Under punctuated models, cladogenesis rates would also affect the likelihoods of character change rates and divergence times simply because even if a rate-of-change parameter remains constant, then the expected amount of change over two different branches with the same duration t will be different if those two branches span intervals with different cladogenesis rates. Innumerable paleobiological studies suggest that cladogenesis rates vary substantially over time (e.g., Foote et al. 2018; Foote 2023; Foote et al. 2024). It is already standard practice for phylogenetic studies to consider a range of possible models of character evolution and diversification simply because we cannot know in advance which set of assumptions maximizes the probability of our observed data or our possible inferences (see, e.g., Wright et al. 2021). Thus, even if speciation models are not a focus of a study that we are conducting, then speciation models might still represent an important nuisance parameter that we need to accommodate to test seemingly unrelated macroevolutionary hypotheses.

We have three goals in this paper. First, we shall elaborate upon a punctuated analog to the Mk model (Lewis 2001) that is commonly used in phylogenetic studies (see also Wagner and Marcot 2010). We will also present means of applying this to existing Bayesian phylogenetic packages such as RevBayes (Höhna et al. 2016). Second, we will then apply this approach to Ordovician strophomenoid brachiopods (Congreve et al. 2015), which represents a clade displaying an early burst of disparity that coincides with elevated rates of cladogenesis. We will use this example to examine the possible implications of punctuated change models when testing “intrinsic constraint” vs. “empty ecospace” explanations for early bursts (Congreve et al. 2021). Third and finally, we will discuss how future alterations of phylogenetic methods might take into account the stasis component of punctuated equilibrium.

## Methods

### Continuous-Time vs Punctuated Change in Phylogenetic Models

Macroevolutionary theory and systematic methods often represent two sides of the same coin: most (if not all) conflicting ideas about ways in which organisms evolve predict different relationships between phylogeny and observable data. Models of character evolution exemplify this. The most common method for anatomical data is Lewis’ (2001) Mk model. Here, the likelihood of a specific combination of relationships, diversification times and rate of change is the Poisson probability of net stasis or ultimately transitioning to another state is given by:

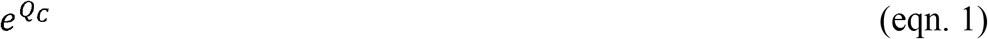

where *Q*c in its simplest form is a transition matrix:

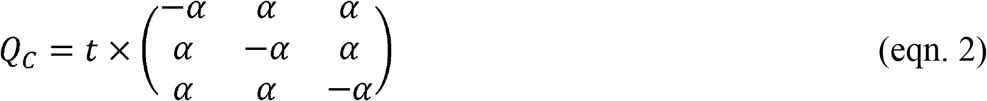

with *t* being the amount of time over which a character might change (e.g., the span between a divergence time and the first appearance of a taxon), α is the instantaneous (Poisson) rate of change, and α*t* therefore is the expected amount of change. (It is unnecessary to place t outside of the matrix and using αt or -αt in each cell will yield the same results; we use this form because phylogenetic analyses vary *t* and α independently, and because the scripts for performing these analyses require separating α and *t*.) Following matrix exponentiation, the probability of net stasis is 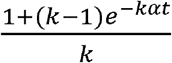 and the probability of any transition is 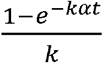 where *k* is the number of character states (e.g., 3 in equation 2; Lewis 2001). Note that the probability of net stasis is not the Poisson probability of no change (*e*^-α*t*^) but instead is the probability of starting at State X and ending at State X. This is because ‘no change’ can be accomplished in two ways: because of truly no character change or because 2+ characters changes ultimately led to reversal. Thus, for a binary character, the probability of net stasis is the Poisson probability of zero changes plus the summed probabilities of all even numbers of change. Similarly, the probability of “ultimate” transition for a binary character is the summed probability of all odd numbers of changes. When there are 3+ states, these probabilities are weighted by the probability that (say) 3 changes will lead back to the original state or from State X to State Y. More complex variants of the Mk model exist that allow for biased state transitions (including driven trends *sensu* McShea 1994), variation in probabilities of state transitions over phylogeny (Nylander et al. 2004; Wright et al. 2016) and correlated character change (e.g., Billet and Bardin 2018). However, these variants all assume that anatomical change can happen at any point in time rather than being concentrated in speciation events.

Because punctuated models posit that change is limited to cladogenetic events, we might expect a punctuated analog to the Mk model to use binomial/multinomial probability. However, this would require that we know the number of cladogenetic events that occur over some duration *t*. If cladogenesis itself is a continuous time process with rate *λ*, then the expected number of events over time *t* is *λt*. If we incorporate the probability of change per branching event is ε, then the expected amount of change over time *t* is simply ε*λt*. This led Lewis (2001: pg. 917) to suggest that the Mk model accommodates punctuated change, making no assumptions about how the change is distributed across a given branch. This will be true only for punctuated anagenetic change in which change is neither continuous over time nor attached to cladogenetic events (Wagner and Marcot 2010). This differs from Eldredge & Gould’s model, which posits that cladogenesis and anatomical change have a common cause. Phylogenetic topologies will imply some minimum number of cladogenetic events, m, which means that expected change is ε(m+*λt*). We can assume m=0 only for a possible anagenetic topology (Fig. 1A). Given a budding cladogenetic topology (Fig. 1B), m=1 for the branch leading to the daughter species and expected change is ε(1+*λt*). Given a bifurcating pattern between two true-sister taxa, there are two possibilities for any one branch (Figs. 1C-D). There must be one branching event, but we usually will have no reason to think that the “necessary” branching event leads to either branch: it might be either a continuation of the ancestral line or it might be the product of cladogenesis. Thus, this prior probability of that the necessary cladogenetic event leads to the branch in question is 0.5 and expected change now is: 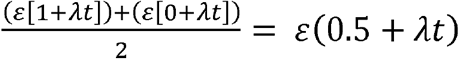. This effectively makes the minimum number of branching events leading to either of two true sister taxa *m*=0.5 and the expected number over branch duration *t* = 0.5+ *λt*. A key difference that emerges is that even if *t*=0, the expected change is greater than zero given any phylogenetic topology in which there must have been at least one cladogenetic event (i.e., *m*=0.5 or 1.0).

**Figure 1.**
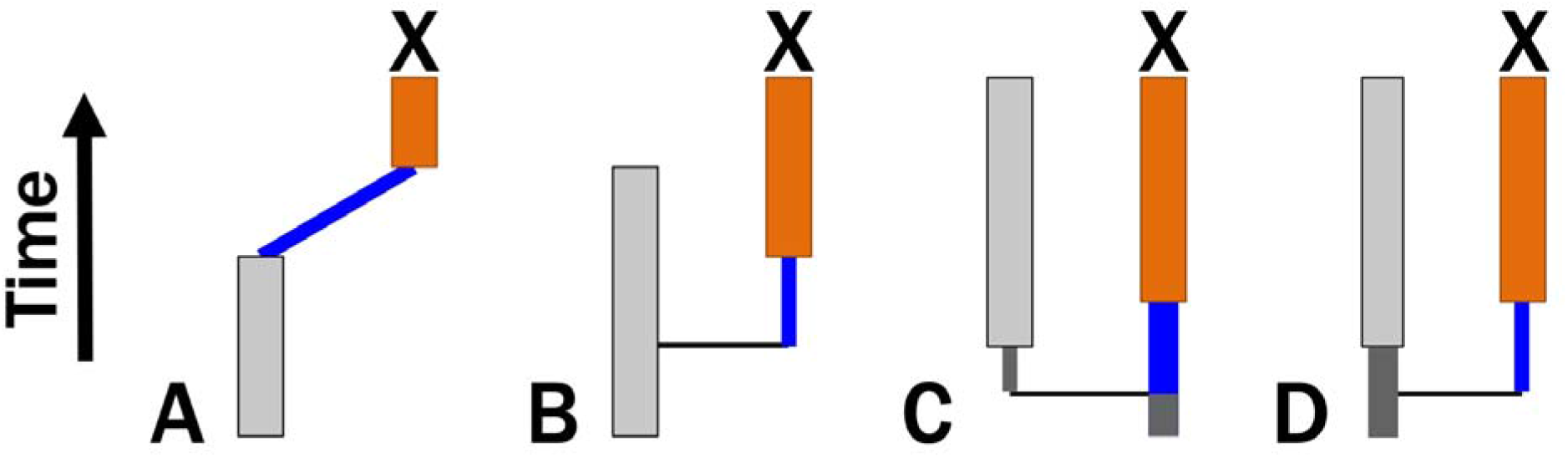
Four possible cladogenetic histories leading to taxon X. Thin black vertical lines denote necessary cladogenesis. Blue lines denote time over which continuous change and/or additional cladogenetic events might occur. A) X is the descendant of an observed species that last appears before X first appears. Here, there might be no cladogenetic events (*m*=0). B) X is the descendant of a species that co-occurs with X. Here, there must be at least one cladogenetic events (*m*=1). C & D) Bifurcation, where neither X nor its sister-taxon are ancestral to one another. There must be one cladogenetic event; however, X might be the continuation of the original ancestor and thus need not have any cladogenetic events separating it from its common ancestor with X’s sister taxon (C). Alternatively, X represents the line derived from the cladogenetic event. Assuming both possibilities to be equiprobable, here *m*=0.5.

Following Wagner and Marcot (2010), a “true” punctuated equilibrium analog to the Mk model in its simplest form is given by:

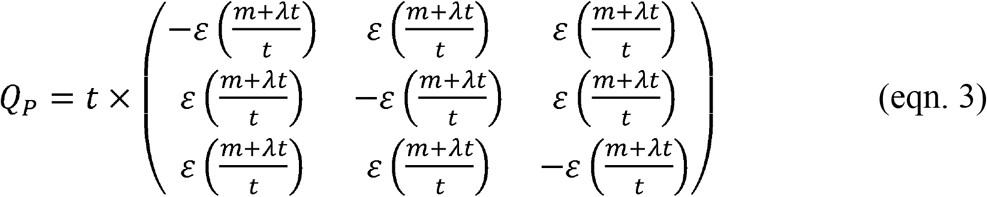

with the probability of net stasis is 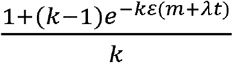 and the probability of “ultimate” transition is 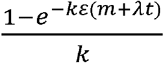. We would accommodate more complex models of character state transitions such as those outline above by varying ε in the same way that those models vary α.

### Cladogenesis rates

Two issues immediately arise from tying expected character change to rates of cladogenesis (*λ*). One is whether allowing two parameters (*λ* and ε) instead of just one (α) to generate the likelihoods of possible evolutionary histories might unduly bias tests towards favoring punctuated equilibrium models. Another is how we go about including *λ* in phylogenetic analyses. Because the latter informs on the former, we will first review how Bayesian analyses incorporate diversification and sampling rates.

Bayesian phylogenetic analyses such as the Fossilized-Birth-Death (FBD) model (e.g., Heath et al. 2014) assess the prior probability of branch durations on phylogenetic topologies based on rates of cladogenesis (*λ*), extinction (μ) and sampling (Ψ) (Fig. 2). Ψ alone affects the probability that we fail to sample either an earlier member of a lineage or some direct ancestor to one or more sampled taxa over time *t* (Wagner 1995a; Foote 1996a). *λ* affects the probability that 0…N cladogenetic events will happen over that time, and the combination of *λ*, μ and Ψ affect the probability that we would fail to sample any descendants of each cladogenetic event (Foote et al. 1999; Stadler 2010; Bapst 2013; Didier et al. 2017). Thus, *λ* has a major effect on the posterior probability of a tree regardless of the model of character change (Fig. 2A) and using *λ* to co-determine expected character change does not introduce any new parameters our model (Fig. 2B).

**Figure 2.**
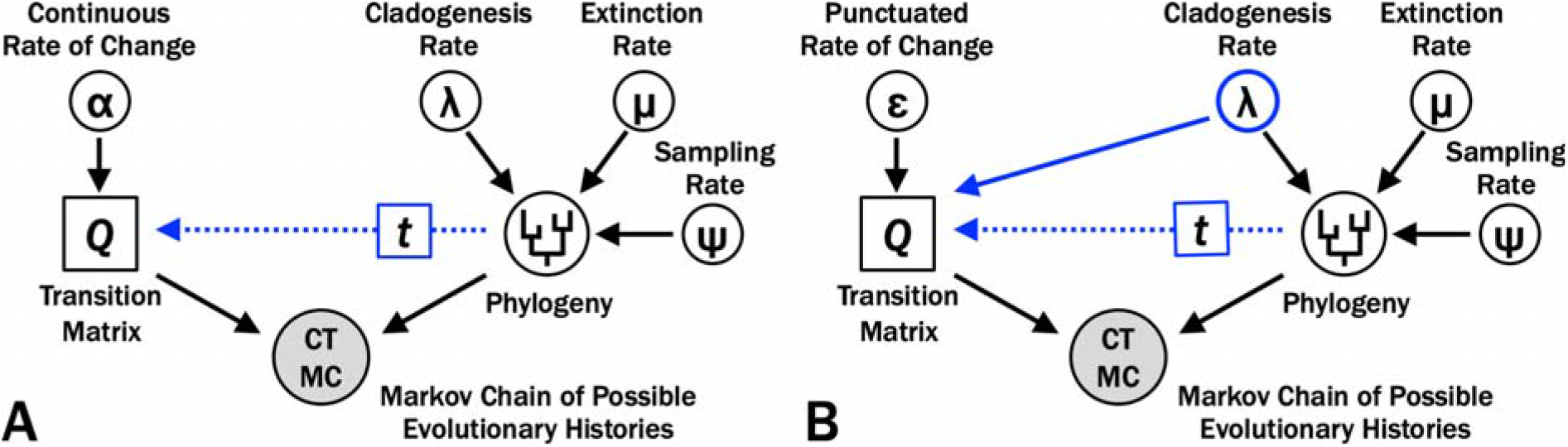
Graphical model for Bayesian analyses of evolutionary histories with varying rates of origination, extinction and sampling over time and constant rates of change among characters and over time. In each graph, the likelihood component is on the left and is evaluated by character data; whereas the prior probability component is on the right and is evaluated by first appearance data. A) Continuous change, where the matrix of transition probabilities (*Q*) reflects the instantaneous rates of change (α) and duration of branches from a particular phylogeny (*t*). B) Punctuated change, where *Q* reflects the probability of change per branching event (ε) and *t*. Although allowing *Q* to reflect two parameters can increase the likelihood component of the posterior probability, this also demands that cladogenesis rate (*λ*) satisfy both first appearance and character data.

As described elsewhere (e.g., Wright 2019; Wright et al. 2021; Wright et al. 2022), we can make numerous modifications to the basic models illustrated in Figure 2 that relax assumptions about homogeneity of rates. We would make identical modifications to both the continuous-time and punctuated models in Figure 2 in order to allow for rate variation among characters (e.g., Gamma or lognormal vs. invariant rates; e.g., Yang 1994; Harrison and Larsson 2015) or over the phylogeny (e.g., relaxed vs. strict clocks; e.g., Sanderson 1997; Drummond et al. 2006). “Skyline models” (e.g., Stadler 2011; Stadler et al. 2013; Warnock et al. 2020) allow for the temporal variation in diversification (or sampling) rates of the sort that innumerable paleobiology studies document (e.g., Krug and Jablonski 2012; Alroy 2015; Foote et al. 2018; Lowery and Fraass 2019; Foote 2023; Foote et al. 2024). Under Skyline FBD models, an X million year branch duration in an interval with high *λ* usually will have a lower probability than will an X million year duration in an interval with low *λ* (see Wagner 2019). This is where the continuous-time and punctuated models might yield very different results, particularly if we are using “strict clock” models for both α and ε. Variation in *λ* over time will allow shifts in expected amounts of change per million years even under constant ε. However, *λ* is simultaneously evaluated by the prior probabilities of branch durations: as *λ* increases, the probability of shorter branches increases and as *λ* decreases, the probability of shorter branches increases (see, e.g., Wagner 2019). Thus, we cannot freely vary *λ* to maximize how well constant (“strict clock”) ε maximizes the probability of the character data. Instead, the punctuated model should result in high posterior probabilities relative to the continuous-time model *only if increasing or decreasing origination rates to improve likelihood given character data also improves prior probabilities given stratigraphic data*.

### Data and Analyses

We examine 37 Ordovician genera from the brachiopod superfamily Strophomenoidea based on character data published by Congreve et al. (2015). Strophomenoids appear during the Middle Ordovician and are one of the two superfamilies (along with the Plectambonitoidea) that contribute most to the diversification of “articulate” brachiopods during the Great Ordovician Biodiversification Event (Harper et al. 2013). Congreve et al. (2021) show that the clade displays a classic “early burst” of morphological disparity and that this coincides with elevated rates of change per million years given phylogenies consistent with the cladogram published by Congreve et al. (2015). However, those elevated rates coincide with similar peaks in origination rates, leading Congreve et al. (2021) to suggest that the early burst might reflect elevated frequencies of punctuated change rather than changes in the basic evolvability strophomenoid shells. The former interpretation is more in keeping with predictions of “empty ecosytems” driving morphological change rather than “relaxed intrinsic constraints” (see, e.g., Valentine 1969, 1980) and thus highlights the potential importance of being able to distinguish between continuous-time and punctuated character change models when addressing other macroevolutionary issues.

We conduct analyses in RevBayes (Höhna et al. 2016). The following aspects of the punctuated and continuous-time analyses are identical for both analyses. Both analyses assume lognormal rate variation among characters, as prior analyses indicate that this fits better than Gamma or invariant distributions (Wagner 2012, Harrison and Larsson 2015). Both used a 12-bin Skyline model, with chronostratigraphic partitions reflecting the Ordovician stage-slices proposed by Bergström et al. (2009) and the Ordovician timescale given in Gradstein et al. (2020), although after lumping some of the relatively brief stage-slices (see Congreve et al. 2021). We seeded sampling (Ψ) and origination (*λ*) rates based on birth□death□sampling analyses published in Congreve et al. (2021) but allowed to vary over the analyses. Following Wright et al. (2021), we did not have extinction (μ) vary independently of origination; instead, we use a separate turnover parameter that follows a lognormal distribution matching substage variation in turnover (μ/*λ*) shown in empirical studies. Finally, the upper and lower bounds of the first appearance times for the analyzed taxa reflect the earliest and latest possible dates of the oldest species in each genus based on data in the Paleobiology Database (PBDB). However, instead of the dates given by the PBDB, we use an extrinsic database (see Congreve et al. 2021) that restricted possible first appearances to conodont or graptolite zones, with the ages of those zone again derived from Gradstein et al. (2020). We could not implement the complex compound Beta distributions for possible first appearance dates used in Congreve et al. (2021); therefore, the analyses assumed a uniform distribution of possible ages within those upper and lower bounds (see Barido-Sottani et al. 2019).

For the purposes of this study, we use invariant rates (= strict clocks) for both continuous□time (α) and punctuated (ε) rates of change. The sole difference in the two sets of analyses is that in the punctuated analysis, the rate of character change was set by ε, the “local” cladogenesis rate (*λ*_i_) and the minimum branching events (*m*). For branches spanning 2+ stage-slices, the expected change from cladogenesis in addition to *m* is given by:

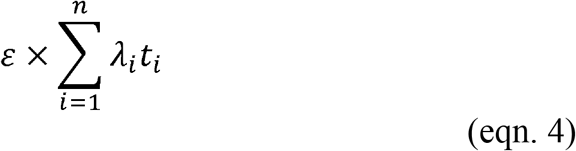

with ∑*t*_i_ equaling the branch duration. We accomplished this by: 1) dividing each branch into 100-kyr intervals; 2) referring to the origin time of each branch on each particular phylogeny; and, 3) setting the rate of change to ε*λ*_i_ where *λ*_i_ is the origination rate during that 100-kyr time slice (see supplementary material). We set *m* to 0.5 for all branches because this was a genus-level analysis and we had no strong intuition of which branches the cladogenetic events should sit on. Even if a particular phylogeny has: 1) one genus as ancestral to another; and, 2) both genera co-existing, then it is possible that the branching event allowing 2+ lineages to co-exist happened within the paraphyletic genus while the “derived” genus represents greater amounts of continuous anagenetic change. This neglects the possibility that an ancestral genus might disappear prior to the first appearance of the derived genus. Unfortunately, Bayesian analyses currently have no way to accommodate last appearance data. Fortunately, initial analyses (e.g., Congreve et al. 2015) do not suggest this happened within the clade and most of the genera have wide ranges with apparent close relatives usually co-existing. As we will discuss below, this is an area that requires further development.

We set up each of our models in RevBayes 1.2.1 (Höhna et al. 2016). We ran the models until convergence, as assessed in the software tracer (about 1 million generations per estimation). RevBayes returns both trees and parameter estimates in a tab-separated format. We then processed results in the R programming environment (R Core Team 2018).

## Results

### Relative Success of Continuous-Time and Punctuated Models

Evolutionary histories of strophomenoid brachiopods with punctuated change are much more probable than are those with continuous change (Fig. 3A). In general, the most probable histories assuming continuous change are only slightly better than are the least probable histories assuming punctuated change. The difference in summed posteriors is large, with the log of the differences >48; this resulting Bayes Factor indicates very strong support for the punctuated model (Kass and Raftery 1995). The distribution in log-posteriors is much more peaked and restricted for the punctuated model than for the continuous-time model. Because the MCMC analyses achieve solid convergence under both models (Supplemental Figure 1), we have no reason to suspect that longer runs would find continuous histories that rival or exceed the best punctuated histories.

**Figure 3.**
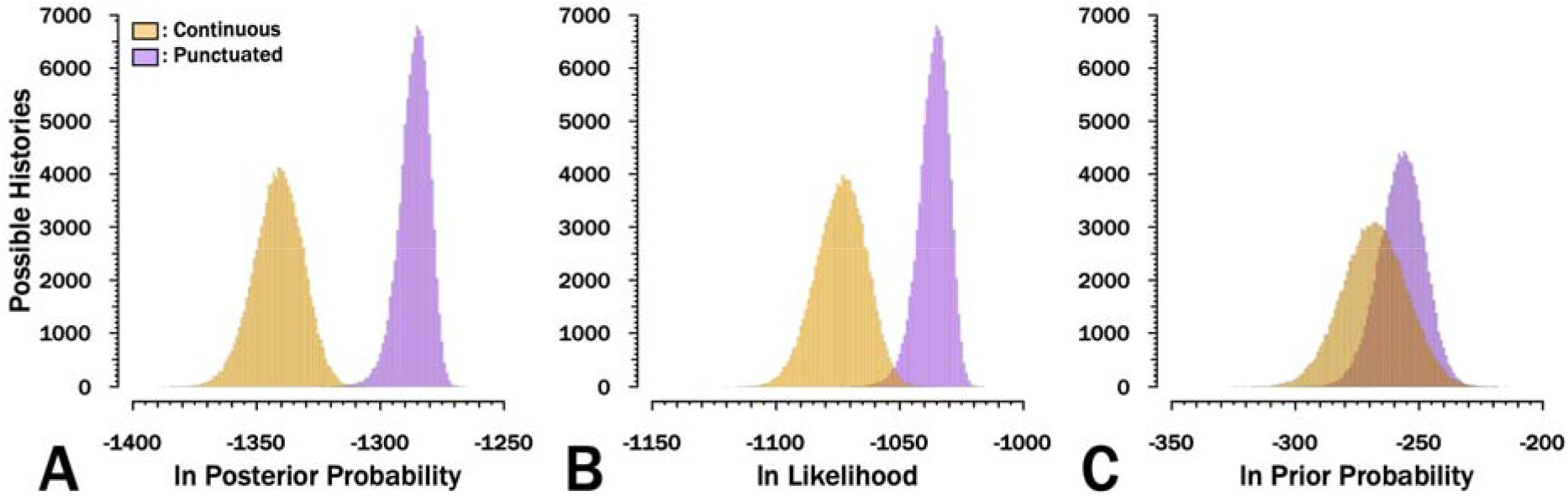
Distribution of results for 105 possible evolutionary histories for strophomenoids saved from 106 total generations. The log likelihoods describe how well either αt or ε(m+*λ*t) predict the character matrix across the phylogeny particular to a given evolutionary history. The log prior probabilities describe well *λ*, μ and Ψ predict the branch durations of the corresponding phylogeny given the range of possible first appearances for the 37 analyzed strophomenoids.

The difference in log-posterior probabilities is driven much more by the differences in log□likelihoods than in differences in log□priors (Figs. 3B vs. 3C). This can be seen in both the differences in values and in how much more constrained and peaked the punctuated posteriors and likelihoods are relative to the continuous posteriors/likelihoods. However, we see the same pattern (albeit less dramatically) for the priors: the prior probabilities of the phylogenetic topologies under punctuated histories tend to be both higher and more concentrated around the central values. This indicates that the punctuated histories do both a better job of predicting the distribution of character state combinations *and* the distributions of first appearances among those character state combinations than do continuous histories. The difference in prior probability is driven predominantly by the prior probabilities of the origination rate, branch durations, and fossil age intervals.

### Apparent Rates of Character Change

Both models employ strict clocks (i.e., the same rates of character change over the entire tree), which means that either α (continuous-time) or ε (punctuated) are the same on all branches. However, heterogeneity in cladogenesis rates (*λ*) over stage□slices results considerable temporal variation in expected rates of change per million years under the punctuated model (Fig. 4). the Dapingian and late Darriwilian (Dw3 of Bergström et al. 2009) both elevate expected change substantially under the punctuated model despite constant ε. Conversely, relatively low cladogenesis rates in the Late Ordovician yield an apparent quiescence in expected amounts of change.

**Figure 4.**
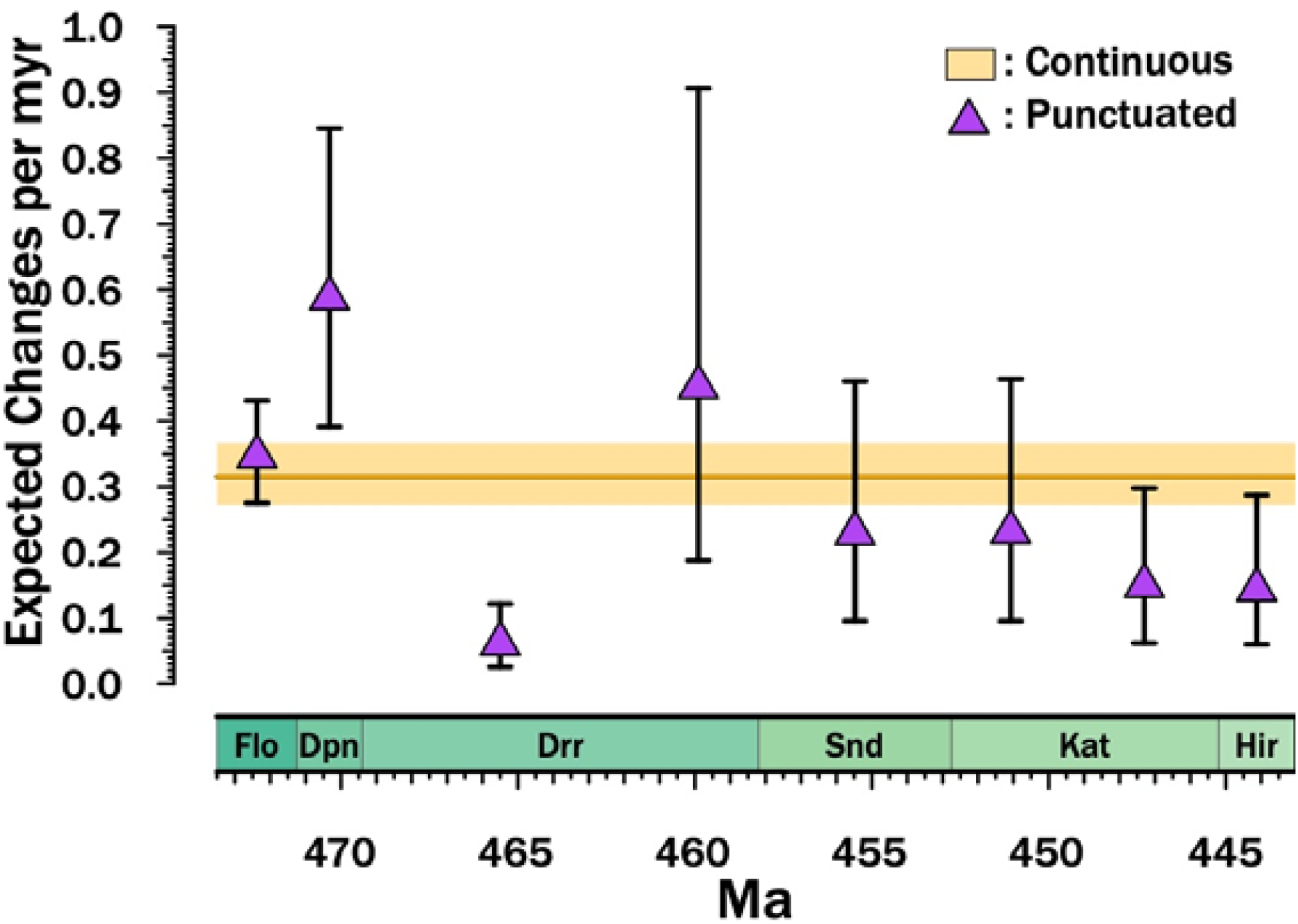
Implied rates of character change per million years. Dark orange line gives the median α from the MCMC runs with the 25^th^ □ 75^th^ percentile distributions shown by the pale orange bar. Purple triangles give the median ε*λ*_i_ from the MCMC runs with brackets encompassing the 25^th^ □ 75^th^ percentile distributions, with *λ*_i_ being the cladogenesis rate for the stage-slice in question. These therefore are directly proportional to the distribution of *λ* within and among stage□slices. For each MCMC iteration, α and ε reflect the median of the lognormal distributions of rate. Finally, note that this excludes the effect of the minimum number of branching events, which means that each branch is expected typically 0.21 changes in the punctuated models in addition to those expected from ε*λ*_i_. Note that most MCMC iterations place the divergence of the analyzed taxa in the earliest Darriwilian (Drr) or latest Dapingian (Dpn); thus, the Darriwilian through Hirnantian points are the ones of primary interest to this study.

### Phylogenetic Relationships

The maximum credibility trees given continuous and punctuated change histories are generally similar, differing more in implied divergence times than in cladistic relationships (Fig. 5). Both trees imply similar patterns of polyphyly among three of the families while suggesting that one (the Rafinesquinidae) is one taxon away from monophyletic.

**Figure 5.**
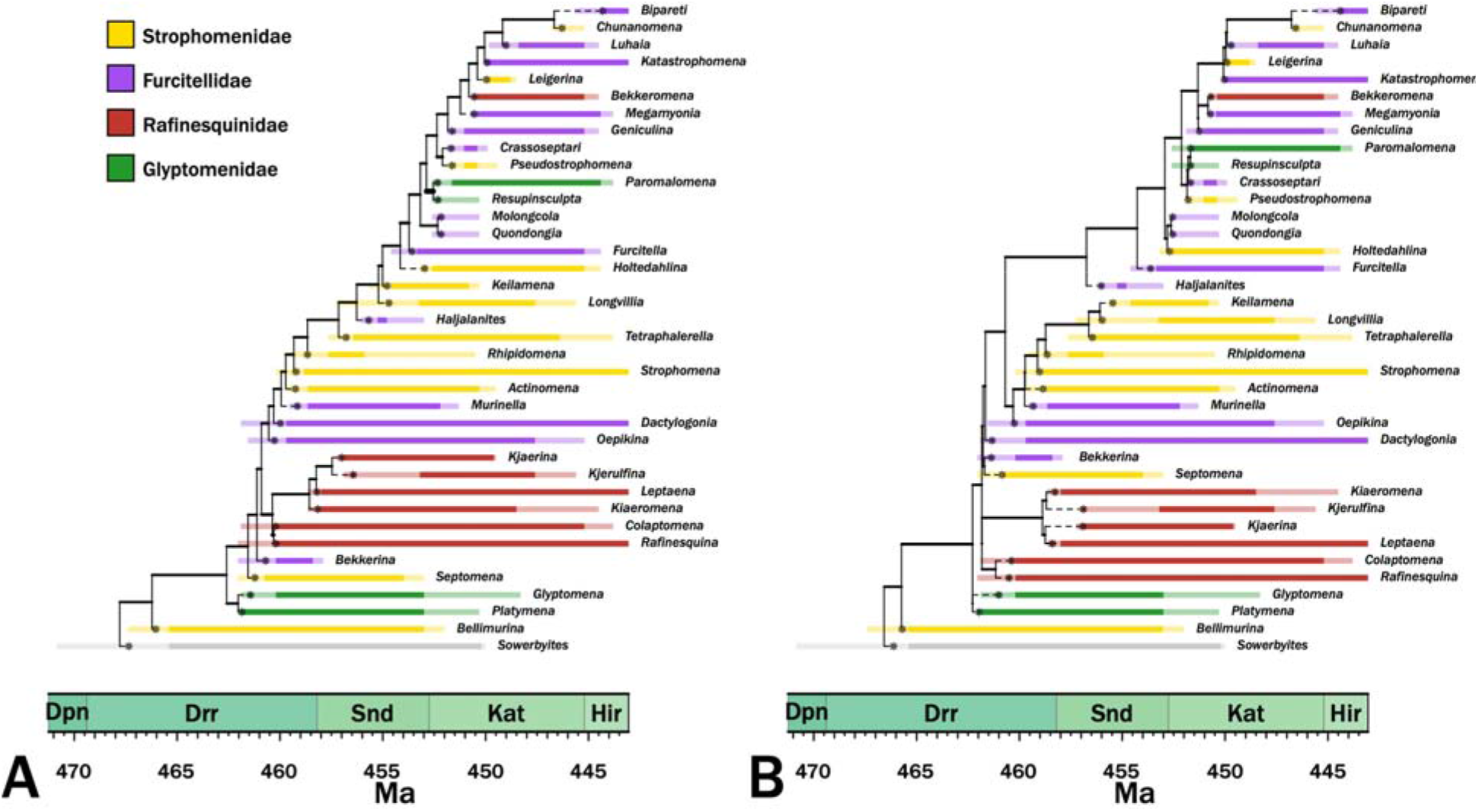
Maximum credibility trees for continuous-time (A) and punctuated (B) models. Colors denote current family assignments. Bold portions denote necessary (range-through) stratigraphic ranges and pale spans denote ranges of possible first or last occurrence ages. Asterisks denote the most probable origins given the analyses.

## Discussion

### Punctuated Character Change among Strophomenoid Brachiopods

Our analyses strongly support the notion that anatomical change within the Ordovician Strophomenoidea is attached to speciation rates and thus consistent with the punctuated equilibrium model. The elevated likelihoods of punctuated histories indicate that the variation in “rates” of change over time generated by temporal variation rates in branching rates do a very good job of predicting anatomical evolution among strophomenoids. It is equally noteworthy that the prior probabilities are generally better given punctuated change histories than given continuous change histories. As we note above, demanding that origination rates predict both branch durations and the distribution of character states among taxa is essentially double jeopardy for the origination rate parameter. In particular, if character change is continuous and cladogenesis rates vary, then intervals of high cladogenesis should generate many short branches with little character change. High rates of cladogenesis elevating the priors under the punctuated model will receive reduced likelihoods due to the lack of change. Conversely, low rates of cladogenesis predicting reduced change along branches will elevate the likelihoods but reduce the priors.

Our results also suggest that the range of data that we can use to assess punctuated change vs. continuous change models is broader than we usually assume. Traditionally, these analyses have focused on species-level studies that relied on contrasting (lack of) change within lineages with apparent change between lineages. Here, we can show at the genus level that explicitly tying rates of change to cladogenesis rates greatly elevates the likelihoods and posterior probabilities of evolutionary histories, and thus is sufficient for assessing a key component of the punctuated equilibrium model.

### Macroevolutionary implications

This study corroborates Congreve’s et al. (2021) suggestion that the elevated rates of change per million years underlying a burst of disparity during the later Darriwilian among early strophomenoid brachiopods was driven at least in part by punctuated change coupled with elevated cladogenesis. This coincides with the peak of the final phase of the Great Ordovician Biodiversification Event in which origination rates were very high for numerous groups (see, e.g., Servais et al. 2010, Stigall 2018, Thomas et al. 2021). These patterns in turn relates to “loose genes” vs. “empty ecospace” notions of what might drive early bursts in disparity (Valentine 1969, 1980). Because “diversity□dependent-” models such as logistic diversification generate elevated cladogenesis rates (*λ*) under empty ecospace scenarios, we expect punctuated change plus rapid radiation to generate high disparity. Of course, this is an incomplete representation of the empty ecospace model, as the original concept focused on relatively lax selection allowing more change than “usual” rather than on elevated cladogenesis rates (e.g., Valentine 1969). Thus, like the “loose gene” models, the empty ecospace model posies (or can posit) elevated α or ε as well as elevated *λ*. Were the point of this paper to fully evaluate strophomenoid evolution during the GOBE, the next step would be to use “relaxed clock” models (see Wright et al. 2021) in which ε varied over time in order to assess whether we do see declining amounts of change per speciation event over time. Moreover, because alterations of modularity or other variations of “loose gene” models might help trigger adaptive radiations (e.g., Wagner and Müller 2002), one would need to partition characters based on inferred ecological roles in order to better distinguish between reduced internal constraints or relaxed ecological restrictions drives elevated ε (e.g., Foote 1994; Wagner 1995b; Ciampaglio 2002, 2004; see, e.g., Wright et al. 2021). However, the primary point for this study is that speciation models are be a critical nuisance parameter when trying to distinguish between basic explanations for rapid increases in morphological disparity. At least in the case of Ordovician strophomenoids, elevated early disparity is at least partially explained solely by elevated punctuated speciation.

### Effects on Inferred Phylogeny

The maximum credibility trees under both continuous change and punctuated change are generally similar, which indicates that the basic patterns are embedded in the character state and stratigraphic range data. We obviously cannot know which tree is more accurate. However, trees from punctuated change histories generally display shallower divergence times and shorter branch durations than do trees from continuous change histories, which is expected if the punctuated change model is generally more accurate than is the continuous change model (Wagner 2000; Wright and Lloyd 2021). Trees from punctuated change history also tend to be somewhat more symmetrical (= “balanced”, i.e., possessing greater proportions of sister taxa that are both clades rather than one clade and one analyzed taxon) than are trees from continuous change histories (Colless imbalance = 0.20 given punctuated; Colless imbalance = 0.35 given continuous-time where 1.0 means a completely “pectinate” or imbalanced tree; Colless 1995), which also is expected if the punctuated model is generally more accurate than is the continuous change model (Huelsenbeck and Kirkpatrick 1996). Although the differences between the maximum credibility trees (and thus the preponderance of the probable trees used to generate those maximum credibility trees) are not great, the fact that they exist suggests that punctuated vs. continuous change represents yet another set of parameters that paleophylogeneticists should consider when assessing phylogeny for its own sake (see Warnock and Wright 2020; Wright et al. 2021).

### Future Developments: the importance of stratigraphic ranges beyond first occurrences

One obvious “flaw” with our approach is that it focuses solely on what happens “between” taxa on a phylogeny by basically treating each taxon as existing in a single point in time somewhere between the lower and upper bound of its first appearance. However, it is quite common for marine invertebrate species to have substantial durations that range through multiple chronostratigraphic boundaries. This in itself has been critical to understanding punctuated equilibrium as it indicates both stasis and situations where one or more long-ranging species gives rise to 1+ daughter species while still persisting (Eldredge 1971). Warnock et al. (2020) lay the groundwork for taking into account stratigraphic ranges beyond the first occurrence in an effort to improve the estimates of extinction and sampling rates for the prior probability of phylogenies. Here, we will outline two other ways in which this should be done to better assess continuous change vs. punctuated change models.

As we note above, the ability of our approach to recognize punctuated change without needing to examine stasis within lineages should expand our ability to test for punctuated character change. The flip-side to this is that being able to modify our tests to account for potential stasis within lineages should greatly expand the power of the tests that we propose. This will be particularly true for species-level analyses where the punctuated equilibrium model predicts that a morphospecies duration reflects only extinction rates (μ) whereas-the continuous change model predicts that a morphospecies duration reflects both μ and the continuous rate of change (α). The probability of observed stasis for one character within a lineage with N finds is:

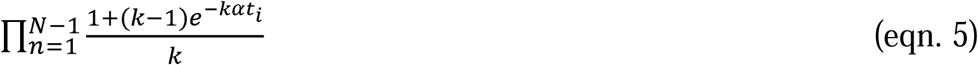

where *t*_i_ is the time that elapses between find *n* and find *n*+1 and 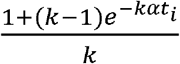 is the probability of net stasis over *t*_i_ given the number of states and α. This introduces an obvious difficulty: just as we know only the upper and lower bounds of possible ages for the first appearance, we will have only upper and lower bounds on the ages of subsequent occurrences. Moreover, even if we have detailed biochronological ordinations of a clade’s species (see, e.g., Guex and Davaud 1984; Alroy 1994; Sadler et al. 2003), then we often will still have numerous finds that cannot be ordered relative to others, resulting in numerous possible combinations of finds. One “shortcut” that is conservative with respect to the continuous change model is to use only the first and last appearances, and thus set *t*_i_ to the duration of the lineage (Wagner and Marcot 2010). This will mean including upper and lower bounds of last appearances and varying both over MCMC searches. By adding this to the likelihood component of Bayesian phylogenetic analyses, the stratigraphic ranges of individual species offer a test of α in addition to the distributions of branch durations and character state combinations.

A directly related issue to stasis and prolonged stratigraphic ranges / temporal durations is the possibility of budding cladogenesis. Indeed, it is often forgotten that the study initiating the punctuated equilibrium debate was a “tree-based” study in which phylogenetic reconstruction of Devonian trilobites suggested that ancestral species survived extinction rates and sometimes generated what we now call polytomies on cladistic depictions of relationships (Eldredge 1971). The maximum credibility trees given either punctuated or continuous change models (Fig. 5) both include the same improbable pattern: numerous long-ranging genera are not reconstructed as “ancestral” (paraphyletic) relative to any other genera (see, e.g., Foote 1996a; Fig. 1C-D). The same issue would be true if this were a species□level phylogeny with numerous long□ranging genera: just as α predicts character change over a species’ lifetime, *λ* predicts that the lineage should be just as prone to giving rise to new species as are the unsampled ancestral lineages linking species on a phylogeny (Wagner and Erwin 1995). There are two issues here. One, because current programs such as RevBayes do not include information about stratigraphic ranges, the MCMC searches could not consider that (say) *Strophomena* was still present when *Rhipidomena* or *Tetraphalerella* appeared, and thus could not consider potentially more probable phylogenies in which one or both taxa “bud” directly from *Strophomena*. Two, the longer a lineage (or group of closely related lineages) persists, the more sampled descendants we expect them to generate. This means that the prior probability of a polytomy with (say) *Strophomena* linked as an ancestor to 2+ descendant species should increase as durations increase (see, e.g., Wagner 2019). Trees lacking such polytomies but still including long□lived species should have lower prior probabilities than should trees placing such species at the “node” of polytomies with descendants appearing at different time over the species’ histories. Note that the mode of speciation itself is not important here: under low rates of continuous change, we will expect accidental “stasis” sometimes, and the same expectations would apply to them. However, because we expect species with long durations relative to origination rates to be more common under punctuated models than under continuous change models, accommodating “natural polytomies” should be more important when change was punctuated.

## Conclusions

Punctuated equilibrium presents a model of character evolution that is fundamentally different than the continuous change models currently used in Bayesian and likelihood phylogenetics. In the case of strophomenoid brachiopods from the Ordovician, punctuated character models imply evolutionary histories that are both more probable and more likely than do continuous change models. Moreover, the punctuated model combined with variable diversification rates over time at least partly accounts for an early burst in strophomenoid disparity purely from elevated speciation rates early in the groups history that elevate the expected numbers of changes per million years without changing the expected numbers of changes per speciation event. Thus, concerns about whether speciation and character change are continuous or punctuated transcends theoretical interest in how speciation operates: it is an important nuisance parameter that can affect our conclusions about hypotheses stemming from seemingly unrelated macroevolutionary theory.

## Acknowledgments

We thank the organizers of this volume for the invitation to submit this work. We would also like to thank N. Eldredge, S. J. Gould and our other predecessors who provided enough thoughts outside of existing boxes that we are still working on expanding the box 50 years later. This research was initiated under NSF□Grants NSF EAR 2129628 to PJW and NSF DEB 2045842 and NSF CIBR 2113425 to AMW. This is Paleobiology Database Publication #XXX.

## Competing Interests

The authors declare that they have no competing interests.

## Data Availability Statement

Data available from the Dryad Digital Repository https://doi.org/10.5061/dryad.k0p2ngf8g.

## Supplementary Information

### I. Traces of Continuous-Time and Punctuated Models

**Figure S01.**
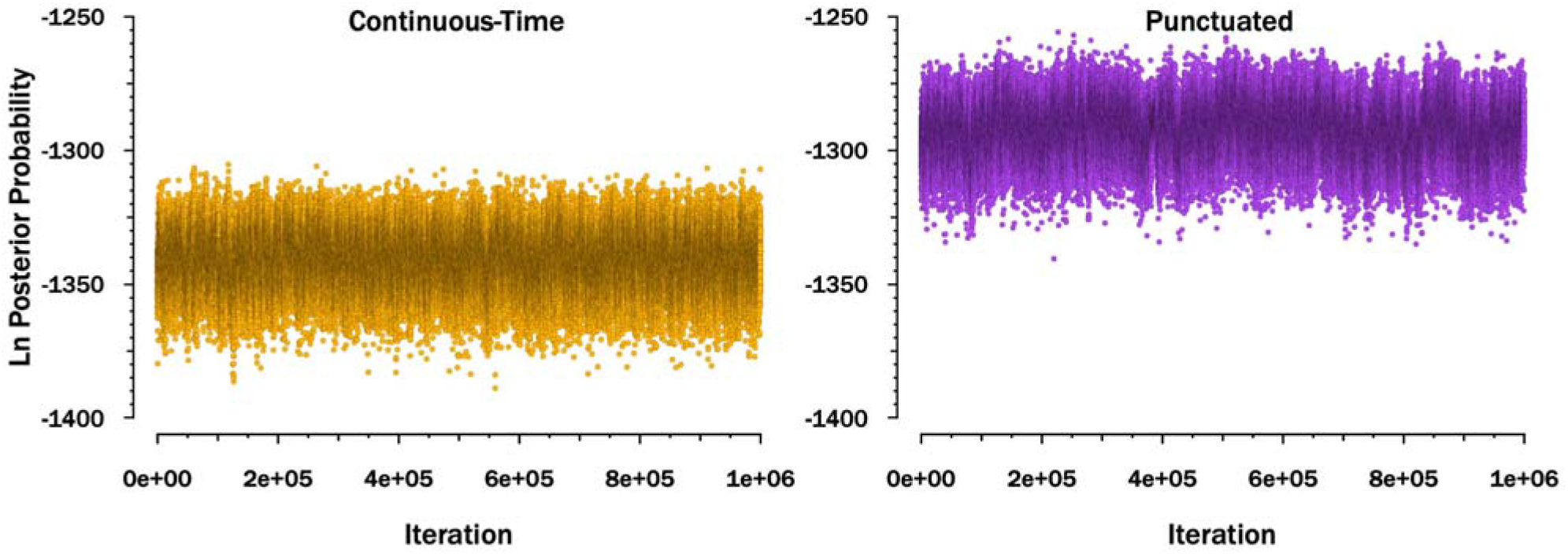
Traces showing the log posterior probabilities of every tenth generation of the Markov Chain Monte Carlo searches.

## Notes

### Competing Interest Statement

The authors have declared no competing interest.

