## Supplemental Material for "Quantitative Models for Distinguishing Punctuated and Continuous-Time Models of Character Evolution and Their Implications for Macroevolutionary Theory"

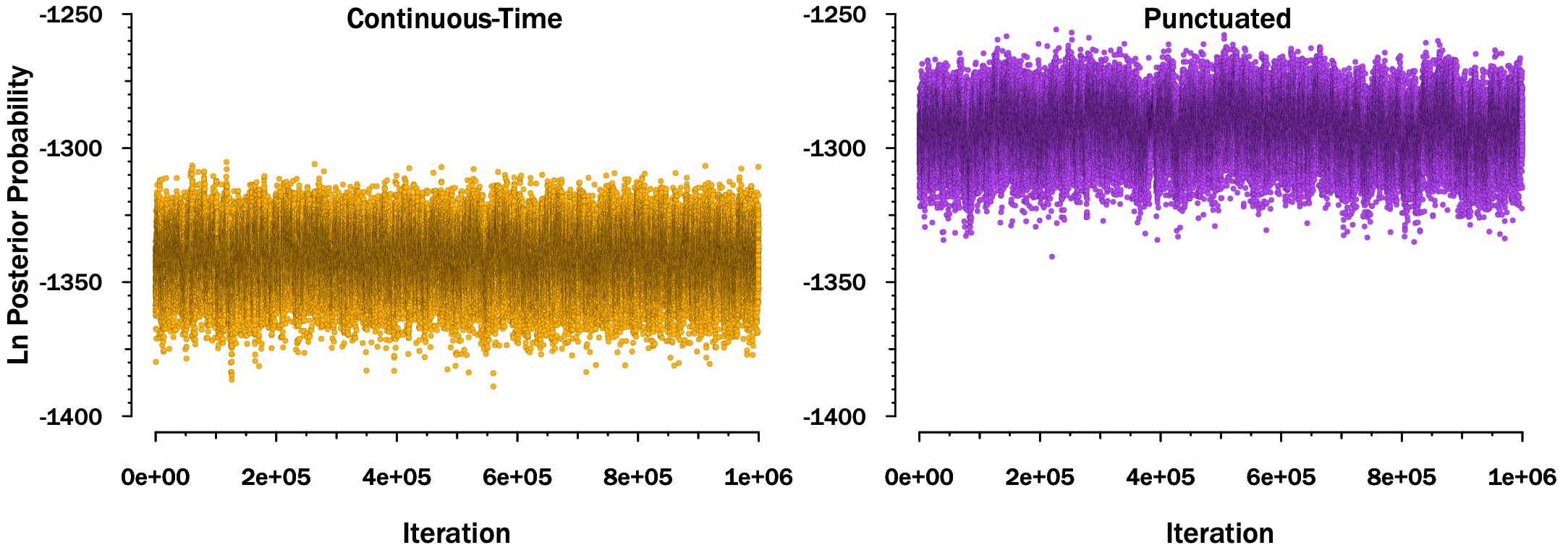

Figure S01 – Traces showing the log posterior probabilities of every tenth generation of the Markov Chain Monte Carlo searches.
